# Targeting and Vaccine Durability Are Key for Population-level Impact and Cost-Effectiveness of a Pox-Protein HIV Vaccine Regimen in South Africa

**DOI:** 10.1101/459909

**Authors:** Christian Selinger, Anna Bershteyn, Dobromir T. Dimitrov, Blythe J.S. Adamson, Paul Revill, Timothy B. Hallett, Andrew N. Phillips, Linda-Gail Bekker, Helen Rees, Glenda Gray

## Abstract

**Background:** RV144 is to date the only HIV vaccine trial to demonstrate efficacy, albeit rapidly waning over time. The HVTN 702 trial is currently evaluating in South Africa a similar vaccine formulation to that of RV144 for subtype C HIV with additional boosters (pox-protein regimen). Using a detailed stochastic individual-based network model of disease transmission calibrated to the HIV epidemic, we investigate population-level impact and maximum cost of an HIV vaccine to remain cost-effective.

**Methods:** Consistent with the original pox-protein regimen, we model a primary series of five vaccinations meeting the goal of 50% cumulative efficacy 24 months after the first dose and include two-yearly boosters that maintain durable efficacy over 10 years. We simulate vaccination programs in South Africa starting in 2027 under various vaccine targeting and HIV treatment and prevention assumptions.

**Results:** Our analysis shows that this partially effective vaccine could prevent, at catchup vaccination with 60% coverage, up to 941,000 (15.6%) new infections between 2027 and 2047 assuming current trends of antiretroviral treatment. An impact of up to 697,000 (11.5%) infections prevented could be achieved by targeting age cohorts of highest incidence. Economic evaluation indicates that, if treatment scale-up was achieved, vaccination could be cost-effective at a total cost of less than $385 and $62 per 10-year series (cost-effectiveness thresholds of $5,691 and $750).

**Conclusions:** While a partially effective, rapidly waning vaccine could help to prevent HIV infections, it will not eliminate HIV as a public health priority in sub-Saharan Africa. Vaccination is expected to be most effective under targeted delivery to age groups of highest HIV incidence. Awaiting results of trial, the introduction of vaccination should go in parallel with continued innovation in HIV prevention, including studies to determine the costs of delivery and feasibility and further research into products with greater efficacy and durability.

## 1. Introduction

With an estimated global prevalence of 36.7 million infected people as of 2015, HIV remains a public health priority in many countries [1]. Despite continued efforts to scale up treatment, resulting in 18.2 million people receiving antiretroviral therapy [2], an estimated 2.1 million people (including 150,000 children) were newly infected in 2015. Reaching the ambitious goal of 90-90-90 by 2020 (90% of people infected with HIV should know their status, with 90% of people diagnosed with HIV infection to be receiving antiretroviral treatment and 90% of people receiving treatment to have viral suppression) would not only diminish a substantial treatment gap but also prevent new infections. However, a recent modeling study including 127 different countries suggests that the majority of countries under consideration (including South Africa) is unlikely to meet the 90-90-90 target [3]. This, and the difficulty in rolling out existing methods of HIV prevention (such as medical male circumcision, and oral pre-exposure prophylaxis) to key populations, highlight the need for a preventative vaccine [4].

October of 2016 marked the launch of the first trial in seven years to test the preventative efficacy of an HIV vaccine in humans. HVTN 702 is a phase 2b/3 trial [5], supported by the Pox-Protein Public-Private Partnership (P5) [6]. It is testing a modified version of the only HIV vaccine to date that has shown evidence of efficacy in humans. In 2009, RV144 showed partial reduction in HIV acquisition among community-based, predominantly heterosexual participants in Thailand using a modified intent-to-treat analysis (vaccine efficacy of 31.2% (95% CI: 1.1% - 52.1%) at month 42 after the first vaccination) [7]. Though the efficacy of the vaccine appeared to be greatest shortly after the last dose and then waned rapidly, a recent follow-up study [8, 9] in which a subset of RV144 participants was re-vaccinated up to six years after enrollment reported immune memory responses two weeks after re-vaccination, offering hope that an extended immunization schedule could potentially increase vaccine durability.

The ongoing HVTN 702 study is a multi-site, randomized, double-blinded, placebo-controlled clinical trial designed to test a regimen adapted to HIV Clade C exposed populations in South Africa. In addition to the six-month series tested in RV144, its original protocol comprises a booster dose at month 12, as well as a change in adjuvant (from alum to MF59) to the gp120 protein component of the vaccine, and a recently amended 18-month booster dose. Preliminary results of HVTN 100 [10], a small-scale clinical trial evaluating the ALVAC-HIV (vCP2438) + Bivalent Subtype C gp120/MF59 (short: ALVAC-HIV-C+gp120) vaccine, suggest good safety, tolerability, and immunogenicity of the modified product and regimen.

If this or an improved pox-protein HIV vaccine regimen is proven to be sufficiently efficacious and licensable, timely and efficient scale-up will require an evidence-based vaccine access plan that defines expectations and outlines commitments essential to making a licensed vaccine available to priority populations in South Africa and potentially beyond. To inform the development of an access plan, the P5 Global Access Committee (short: GAC, comprised of representatives from the Bill & Melinda Gates Foundation, National Institutes of Health (DAIDS/NIAID), Sanofi Pasteur, GlaxoSmithKline and the South African Medical Research Council) has engaged in a variety of preparatory analyses-including commissioning the modelling work detailed in this article-to identify the populations that would benefit most from the vaccine. Here, target populations for vaccination were solely defined based on age and sex, although other key populations (e.g. commercial sex workers) could have similar benefits from vaccination should the product prove efficacious.

In South Africa, HIV incidence varies considerably by age and sex. The highest HIV incidence rates are observed in young women aged 15-24. In 2012, the estimated incidence in this age group was 2.54 % per year (95% 2.04-3.04%), which is five times the rate of HIV incidence in young men aged 15-24 [11, 12]. Recent results from an observational HIV study in rural KwaZulu-Natal suggest the highest risk of infection among women aged 18 to 28, and men aged 23 to 33 [13, 14]. These epidemic patterns imply that prioritizing high levels of vaccination among young women and a slightly older age group of men would most efficiently reduce new HIV infections.

However, determining the optimal use of an HIV vaccine based on age patterns of incidence has several complications. First, an optimal vaccination schedule should account for any waning efficacy, recommendations for booster dose frequency, and rate of attrition from the recommended series of HIV vaccine doses. Second, a licensed vaccine may take up to a decade before being approved for widespread use in South Africa. In the interim, nationally representative HIV epidemiologic patterns are likely to change from current data, which dates back to 2012. Forecasting changes in incidence patterns is needed to predict the likely impact of implementation scenarios under consideration and design a relevant and effective scale-up strategy. Finally, the patterns of HIV transmission must be accounted for in order to fully capture the potential population-level impact of a vaccine. Effective vaccination will prevent HIV infections not only in the direct recipients, but also in their sexual partners within the contact network.

Several modeling studies [3, 15, 16, 17, 18, 19, 20, 21, 22, 23, 24] have estimated impact and cost-effectiveness of an RV144-like vaccine, but all have assumed either a constant level of efficacy or exponentially waning immunity after a single course of vaccination. In addition to bridging these gaps, the present work offers additional improvements such as application of an age-structured HIV network model, more realistic date of vaccine introduction, and complex booster schedules. Here, we estimate the impact of a 10-year vaccine regimen (primary series and boosters) within a 20-year vaccination program on the HIV epidemic in South Africa using an individual-based network model of HIV transmission structured by age, sex and risk. We model the efficacy profile associated with the 5-dose regimen following the original HVTN 702 protocol and include possible booster doses following the primary series (beginning at 36 months from the first vaccination). The present analysis evaluates and compares the impact and cost-effectiveness of implementation strategies initiating HIV vaccination based on targeted age groups, coverage goals, booster attrition, roll-out, HIV treatment scale-up, and oral pre-exposure prophylaxis (PrEP) availability.

Results from the present modelling analysis will help to inform ongoing vaccine access planning elements, including priority populations for whom the pox-protein HIV vaccine would be expected to have the greatest and/or most efficient public health impact.

## 2. Methods

We developed an agent-based model of the South African population to forecast HIV infections, disability-adjusted life-years (DALYs), and healthcare costs from a government payer perspective over a 30-year time horizon, from year 2018 to 2047. As compared to a reference case with no HIV vaccine, we evaluate implementation of strategies for initiation of HIV vaccination.

### Model set-up and calibration

We modified EMOD-HIV v2.5, an age-stratified and individual-based network model of HIV of South Africa, to incorporate HIV vaccination according to pox-protein HIV vaccine regimens (such as the regimen currently being tested in HVTN 702). Because EMOD is an individual-based model, interventions such as a time-varying course of vaccine efficacy can be applied to each individual according to his or her own timing of vaccination and adherence to the booster series. This renders the model well suited for a nuanced analysis of the anticipated time-dependent efficacy of the pox-protein HIV vaccine regimen.

To ensure that our analysis reflected the realities of the HIV epidemic and the health system in South Africa, we iteratively engaged South African government, academic and community stakeholders through one-on-one interviews, a vaccine access planning summit and a public health impact modelling workshop during which preliminary results of this work were presented and discussed. This stakeholder engagement process helped us understand perspectives on the future HIV prevention landscape, benefits of and challenges to reaching specific target populations and the economic factors that will influence vaccine access, all of which were incorporated into our analysis and considerations.

The parameters, model input values, sources, projections, and sensitivities of the epidemic projection without vaccine, used as the reference group for comparison, have been described previously [25, 26, 27]. A detailed model description, user tutorials, model installer, and source code are available for download at http://idmod.org/software.

For detailed information on the model set-up, calibration and baseline assumptions on HIV treatment and prevention other than vaccine we refer to the Supplementary Material.

### HIV Vaccine Efficacy

We incorporated a parametric model of time-dependent vaccine efficacy that was hypothesized for the pox-protein regimen based on results from RV144. We included the time series of efficacy associated with each dose administered during the study and possible booster doses beyond the 24-month duration of the first stage of the study. The original pox-protein dosing schedule administered a series of five immunizations over 12 months (the 18-month dose recently amended to the protocol was not modeled here). ALVAC-HIV-C was administered at months 0 and 1, followed by ALVAC-HIV-C+gp120 at months 3 and 6, and ALVAC-HIV-C+gp120 dose was administered at month 12 (supplementary to RV144 schedule). Time-dependent vaccine efficacy was interpreted as a per exposure reduction in the probability of acquisition parameterized by an impulse and exponential decay model.

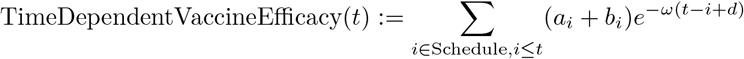

where *a_i_* is the efficacy increase of immunization with ALVAC-HIV-C, *b_i_* is efficacy increase after ALVAC-HIV-C + gp120 immunization, *ω* is the efficacy decay rate per month and *d* is the delay between immunization and initiation of protective effect in months.

Assuming uniformly distributed exposure over a given time span in the trial, we calculated the cumulative vaccine efficacy (corresponding to the efficacy estimate from the trial) as the area under the curve of the instantaneous vaccine efficacy rescaled by the length of the time span.

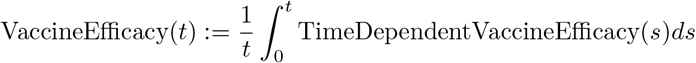

In anticipation of efficacy results for HVTN 702, we modeled time-dependent vaccine efficacy based on results from statistical models [28] for RV144 study outcomes using a point-estimate of 58% shortly after the month 6 vaccination, and cumulative efficacy of 31.2% over 42 months. We adjusted the parameters of the efficacy function such that the cumulative vaccine efficacy over 24 months after the first dose is 50% (corresponding to the goal set by the P5) and obtained values *a_i_* = 0.08, *b_i_* = 0.34, *ω* = 0.065 and *d* = 0.1.

### Booster Schedule and Efficacy

For the purpose of model projections beyond the primary trial endpoint, we also implemented up to four two-yearly boosters starting at month 36 with fixed attrition rates of 0, 20 or 50% per booster to cover a total of 10 years of vaccine efficacy. We assumed booster efficacy to follow the same parameterization as ALVAC-HIV-C + gp120 doses from the primary immunization series during the first 12 months. Booster eligibility depended on having received the primary immunization series or the booster previously. Missing a booster resulted in loss of eligibility for subsequent boosters. Individuals who tested HIV positive were not eligible for future boosters, and we did not add HIV testing to the cost. We assumed that four booster doses after the primary first-year series were necessary to confer one decade of protection. For catch-up vaccination scenarios, booster eligibility was limited by the age range of the vaccinee.

### Vaccination Strategies for Evaluation

Based on consultations with the P5 GAC, we evaluated 30 implementation strategies for HIV vaccine targeting and implementation. Strategies included a wide range of age- and gender-specific targeting scenarios at low (30%), medium (50%) and high (80%) coverage using two distinct roll-out approaches starting in 2027, the earliest realistic date of introduction: Cohort vaccination, aiming for immunization and follow-up boosting of a pre-specified proportion (coverage) of individuals at a particular target age every year over the period between 2027 and 2047 (Table S1), and catch-up vaccination scenarios (Table S2) were implemented for individuals within a pre-specified age range starting in 2027 with a linearly increasing coverage to reach the ramp-up coverage in 2032, followed by cohort-like vaccination from 2032 onwards for individuals aging into the target range at a maintenance coverage which was 20% higher than the ramp-up coverage. For both roll-out scenarios, we assumed no coverage attrition for the primary immunization series. Eligibility for vaccination was assumed to be independent of PrEP usage, and individuals having tested positive for HIV were not eligible for vaccination.

### ART and PrEP Scale-Up Scenarios

To account for uncertainty in predicting the next decade of the HIV epidemic, we varied the scale-up of treatment in terms of ART and oral PrEP coverage and considered three scenarios. In the most pessimistic scenario (Status Quo without PrEP), we stipulated that guideline changes have no impact on ART initiation. We excluded any use of oral PrEP. These assumptions were consistent with the 60% ART coverage under current guidelines assumed in the HIV/TB Investment Case Report for South Africa [29] and reflect to some extent recent pessimistic results on treatment linkage and scale-up in South African settings [3, 30]. A moderately optimistic scenario (‘Status Quo with PrEP’) maintained ART linkage assumptions of ‘Status Quo without PrEP’, but assumed reaching oral PrEP coverage of 30% by 2027 and maintained this level for high risk men and women under 30 years of age. In the most optimistic scenario (‘Fast Track with PrEP’), we kept the same PrEP coverage as in scenario ‘Status Quo with PrEP’, increased testing and linkage to ART and decreased lost-to-follow-up which results in close to 90% ART coverage in 2047. We did not incorporate behavior change into the ‘Fast Track with PrEP ‘scenario, as detailed in the Fast Track UNAIDS goals [31].

### Metrics for population-level health impacts

Epidemic impact was estimated in terms of the number of HIV infections prevented over the period between 2027 and 2047, calculated as differences between the total number of infections given a vaccination strategy and the reference with no HIV vaccine, over the 20-year time period. The fraction of cumulative infections prevented was calculated as the ratio of new infections averted to the number of new infections accumulated between 2027 to 2047 in the reference case (covering thus the period of vaccination). For each implementation policy strategy evaluated, we sampled with replacement from 50 simulation results (corresponding to the 50 most likely parameter sets obtained from the calibration process, see Supplementary Material) to obtain bootstrap mean estimates. The 2.5 and 97.5 percentiles of the sample mean distribution formed the lower and upper bounds of the bootstrapped confidence intervals [32]. This accounts for parameter uncertainty within a neighborhood where likelihood is maximal. We report infections prevented by rounding to thousands.

We also estimated the number needed to vaccinate (NNV) per infection prevented, defined as the ratio of the average number of vaccine regimens distributed to the average number of new infections prevented. The number of vaccine regimens distributed was defined as the number of primary series completed. Health impact was summarized in disability-adjusted life years (DALYs), calculated as the present discounted value of years of healthy life lost to disability and years of life lost to premature death over a 20-year horizon of vaccination [33]. Disability weights for HIV health states differed by CD4-count category (Table 1). DALYs averted were calculated by subtracting cumulative DALYs with vaccine from those without vaccine for each of the three treatment and prevention scale-up scenarios. We measured health outcomes in DALYs to capture changes in both the length and quality of life for individuals in the population. DALYs were discounted 0, 3 or 5% annually.

**Table 1:** Parameters for health impact and economic evaluation analysis

| Healthcare costs, adjusted to 2015 US\$ | | | | |
| --- | --- | --- | --- | --- |
| Cost of vaccine | Wholesale acquisition cost of HIV vaccine product, cost of delivery, implementation, distribution, clinic visits, per regimen |  | unknown |  |
| Cost of PrEP | | | 287 US\$ | |
| Drugs | Average cost per person-year | | 172 US\$ | [16, 29] |
| Visits, Testing, Labs | | | 115 US\$ | |
| HIV care cost | | | 419 US\$ | |
| ART drugs | Average cost per person-year | | 197 US\$ | [16] |
| Labs | | | 90 US\$ | |
| Salaries | | | 66 US\$ | |
| Outpatient, others | | | 66 US\$ | |
| Health outcomes |  |  |  |  |
| Disability Weights for HIV, given CD4-count |  |  |  |  |
| > 350 | Unit disability weights per life year |  | 0.053 | [34, 35, 36] |
| 200 – 349 |  |  | 0.221 |  |
| < 200 |  |  | 0.547 |  |
| Other variables |  |  |  |  |
| CET | | | 5,691 US\$ | South African GDP per capita in 2015 [37] |
| | Cost-effectiveness threshold in 2015 US\$/DALY averted | | 750 US\$ | HIV/TB investment case, Table 54 in [29] |

### Maximum Vaccine Costs

The launch price and implementation cost of an HIV vaccine in South Africa are unknown. Following a value-based pricing framework for medicines [38, 39, 40], a threshold analysis varied the vaccine cost parameter value to identify potential maximum cost-effective HIV vaccine costs (including both wholesale acquisition and implementation cost), as an average across persons receiving different numbers of doses.

Based on strategy-dependent changes in total discounted healthcare costs and DALYs, as compared to the reference with no vaccine, we estimated the maximum vaccination cost level for a vaccination strategy to be cost-effective. This cost level included both the ALVAC and the protein component, as well as implementation cost (e.g. personnel cost of vaccine delivery, facility-level costs, supply-chain management). Since we assumed complete primary series, wasted doses could potentially originate only from missed two-yearly boosters. South Africa does not have an established cost-effectiveness threshold per DALY-averted to determine value for money, although some recent estimates are now available [29, 41, 42]. The cost-effectiveness threshold should represent the opportunity costs of committing scarce health system resources to an intervention [41]. To aid with interpretation, we assumed a threshold of 1x the 2015 gross domestic product (GDP) per capita in South Africa as a cost-effectiveness threshold as well as a threshold of approximately 750 USD per DALY averted, as determined by the South African HIV/TB investment case study group (see Table 54 in [29]).

### HIV Vaccine Price Thresholds

For a given nominal vaccine cost and vaccination strategy, we calculated the net DALY burden relative to the no vaccine reference case at a given cost-effectiveness threshold (CET):

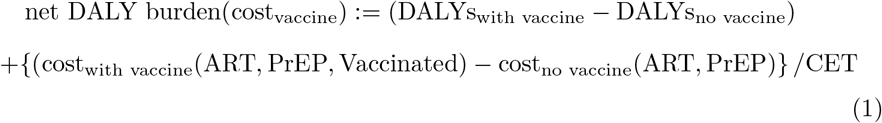

Cost and DALYs, accumulated between 2017 and 2047, were averaged across stochastic replicates, treatment scale-up scenarios and booster attrition levels, and discounted annually at a fixed rate of 0, 3 or 5%. The strategy-specific maximum vaccine cost that remains cost-effective was defined as the maximum regimen cost such that net DALY burden would remain negative, indicating a reduction in population burden of disease (i.e. health gain) from vaccine and improvement in societal welfare. Bootstrapped confidence intervals for vaccine cost were obtained by the same method as described above in the section on population-level impact.

## 3. Results

### Incidence given no HIV vaccine

Varying assumptions about linkage to and drop-out from ART and uptake of oral PrEP between 2016 and the presumed start of vaccination in 2027 results in distinct future ART coverage and incidence estimates. For the scenarios ‘Status Quo without PrEP’, ‘Status Quo with PrEP’ and ‘Fast Track with PrEP’, the average percentage of HIV-diagnosed individuals on ART would be 56, 55 and 85% respectively in 2027, compared to 54% in 2016 (Figure S2). Likewise, for the same scenarios, annual HIV incidence in women aged 15 to 49 is projected to decline from 1.6% in 2016 to 1.21% for ‘Status Quo without PrEP’, 1.17% for ‘Status Quo with PrEP’, and 0.76% for ‘Fast Track with PrEP ‘by the year 2027. During the same time period, annual HIV incidence in men aged 15 to 49 is projected to decline from 0.9% to 0.7%, 0.68% and 0.41% respectively (Figure 1). The gender discrepancy in incidence is maintained, with a male-to-female incidence ratio between 0.57 and 0.54 in 2027 regardless of the scale-up assumptions. For a general population aged 15-49, the average incidence rate in 2027 across all three scale-up and prevention scenarios is projected to be 0.81%.

**Figure 1:**
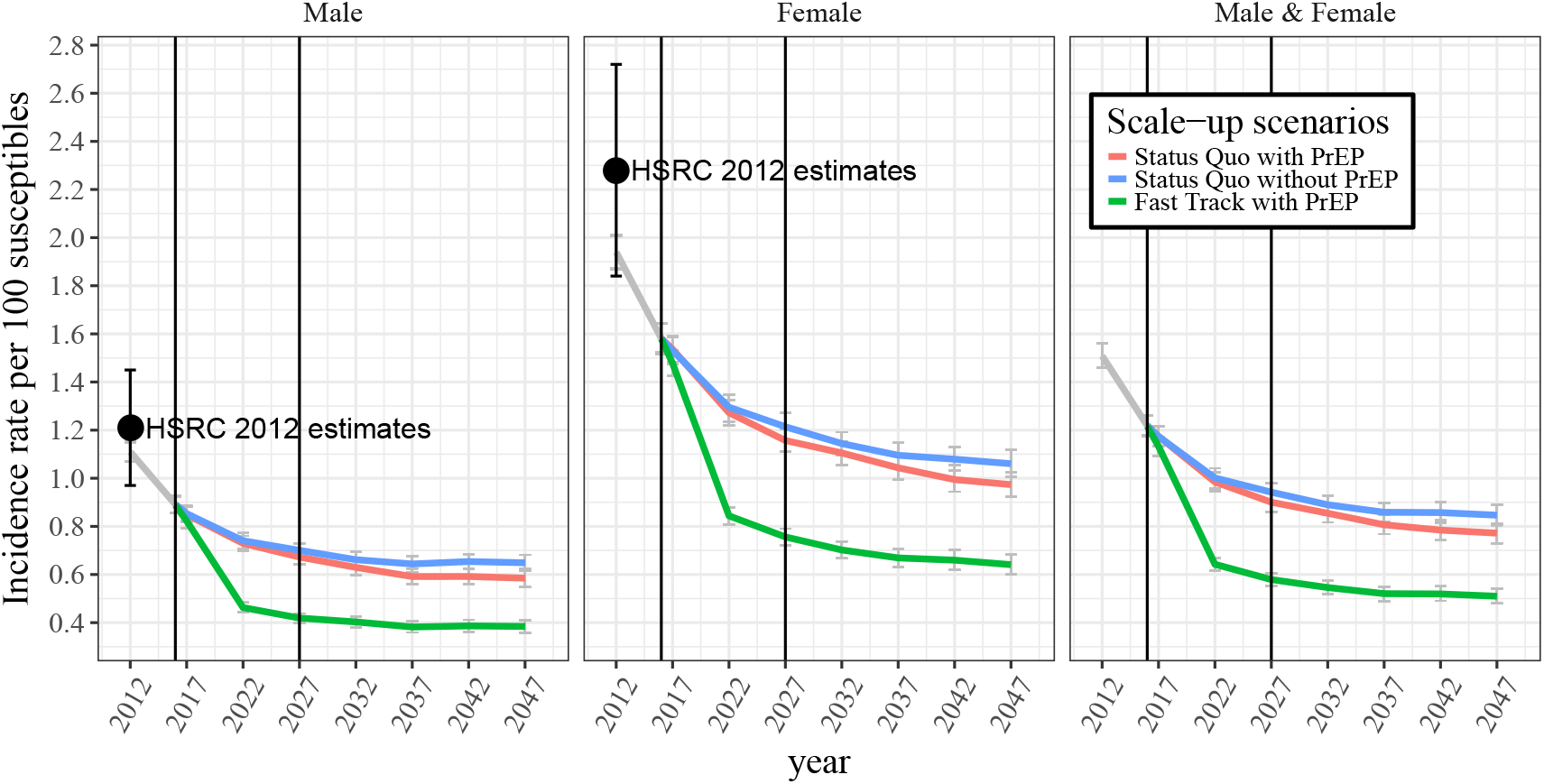
Averaged incidence rates projected under three different treatment and prevention scale-up scenarios starting in 2016, without vaccination: ‘Status Quo without PrEP’ in blue, ‘Status Quo with PrEP’ in red, ‘Fast Track with PrEP’ in green, and the average across all three scale-up scenarios in grey.

Without the vaccine, model projections predict an average of 0.84%, 0.76% and 0.48% annual HIV incidence in a general population aged 15-49 by 2047 for ‘Status Quo without PrEP’, ‘Status Quo with PrEP’ and ‘Fast Track with PrEP’ respectively. This indicates that introduction of oral PrEP for high-risk groups starting in 2016 will have a smaller impact on overall HIV incidence as compared to ART scale-up.

### Time-dependent and cumulative vaccine efficacy

Fitting the impulse and exponential decay model to known time-dependent vaccine efficacy estimates [28] for the first six months of the RV144 regimen, and adjusting the model parameters such that the goal of 50% efficacy at the month 24 endpoint is met, results in a time-dependent efficacy curve (Figure 2, red curve) peaking at 80% shortly after the additional month 12 booster. Exponential waning rates are estimated at 0.065 per month, and with each dose of ALVAC-HIV-C+gp120, incremental efficacy increases by 34%. Two-yearly boosters starting at month 36 will sustain durability with a maximum of 29.6% cumulative efficacy over one decade of vaccination, whereas discontinuing boosters after the first-year series yields a reduced cumulative efficacy of 14.8% (Figure 2, green curve). For the latter case, estimated efficacy of 40% at month 39 coincides with predictions from recently developed statistical approaches using immune correlates of protection [43].

**Figure 2:**
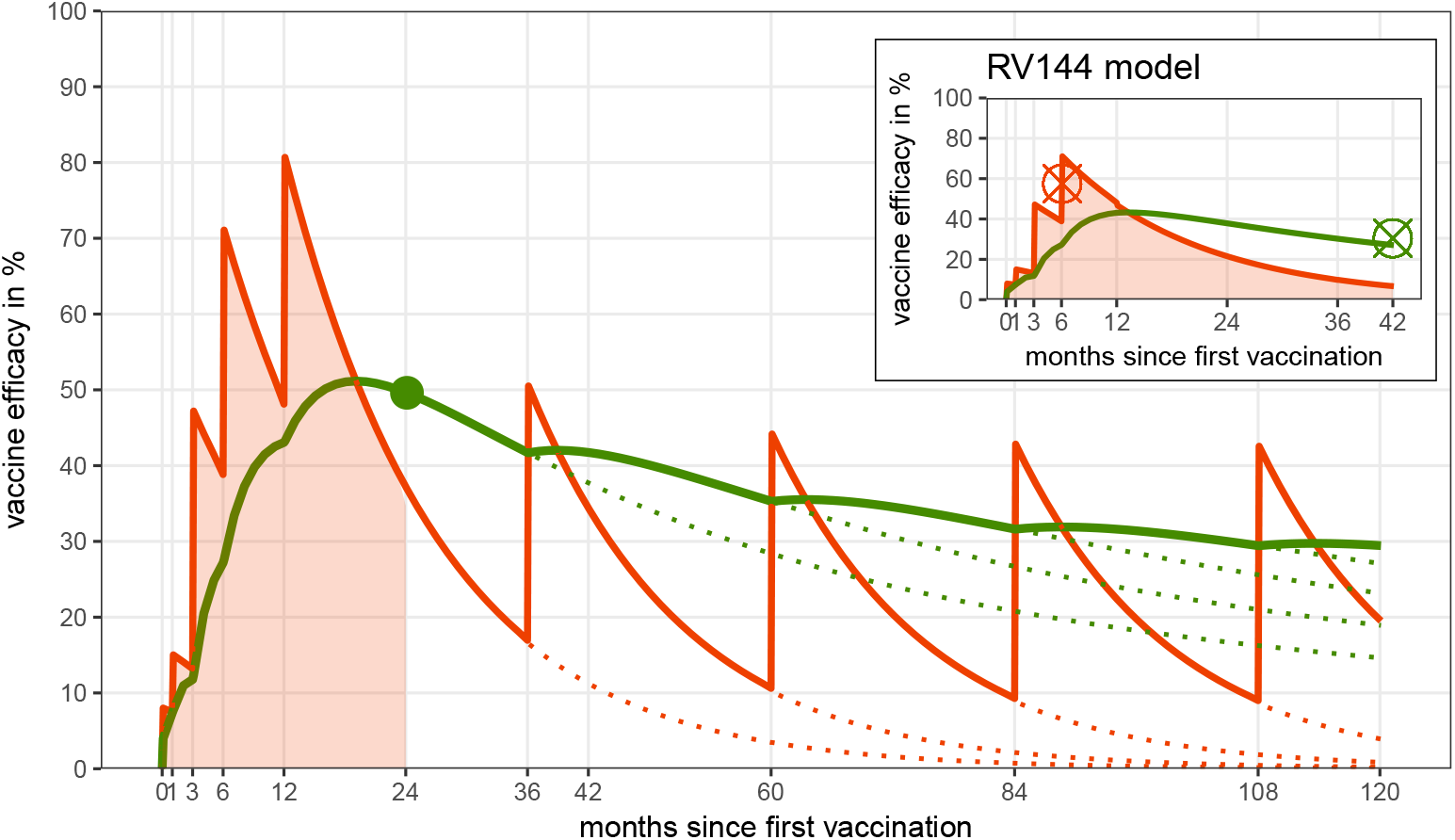
Time-dependent vaccine efficacy (red) is modeled by a parametric impulse and exponential decay model. Cumulative vaccine efficacy at a given endpoint (green) is interpreted as the area under the time-dependent vaccine efficacy curve (shaded red) normalized by the length of the considered time period. We first fit to RV144 point estimates at month 6 and the 3 years endpoint of cumulative efficacy (red and green cross in the small panel). Then we adjusted parameters of the time-dependent vaccine efficacy curve to the P5 regimen schedule such that the goal of 50% efficacy at the 24 month endpoint is met (green point). Dotted lines refer to time-dependent vaccine efficacy with continued booster vaccination after month 24.

### Impact of targeting and durability

To determine vaccination age-targeting of highest impact, we compare cohort vaccination at age 18 to age 15, with or without a 5-year age off-set between men and women (Figure 3) at a relatively high coverage of 80%, averaged across the treatment and prevention scale-up scenarios (see Figure S3 for 50% coverage outcomes). Vaccine eligibility at 15 years of age with full booster retention for one decade prevents between 321, 000 (95% CI: 312, 000 – 329, 000) and 504, 000 (95% CI: 494, 000–516, 000) new infections over twenty years, assuming ‘Fast Track with PrEP’ and ‘Status Quo without PrEP’ respectively. This corresponds to 8.28% (95% CI: 8.11–8.45%) and 8.34% (95% CI: 8.17–8.52%) of new infections prevented respectively. We note that the difference in percent new infection prevented between the two scale-up scenarios is negligible, since the number of new infections in the counter-factual (i.e. without vaccine) for ‘Fast Track with PrEP’ is substantially lower than in the ‘Status Quo without PrEP’ scenario. High booster attrition at 50% would decrease these numbers to 176, 000 (95% CI: 168, 000 – 185, 000) and 309, 000 (95% CI: 296, 000 – 320, 000) respectively.

**Figure 3:**
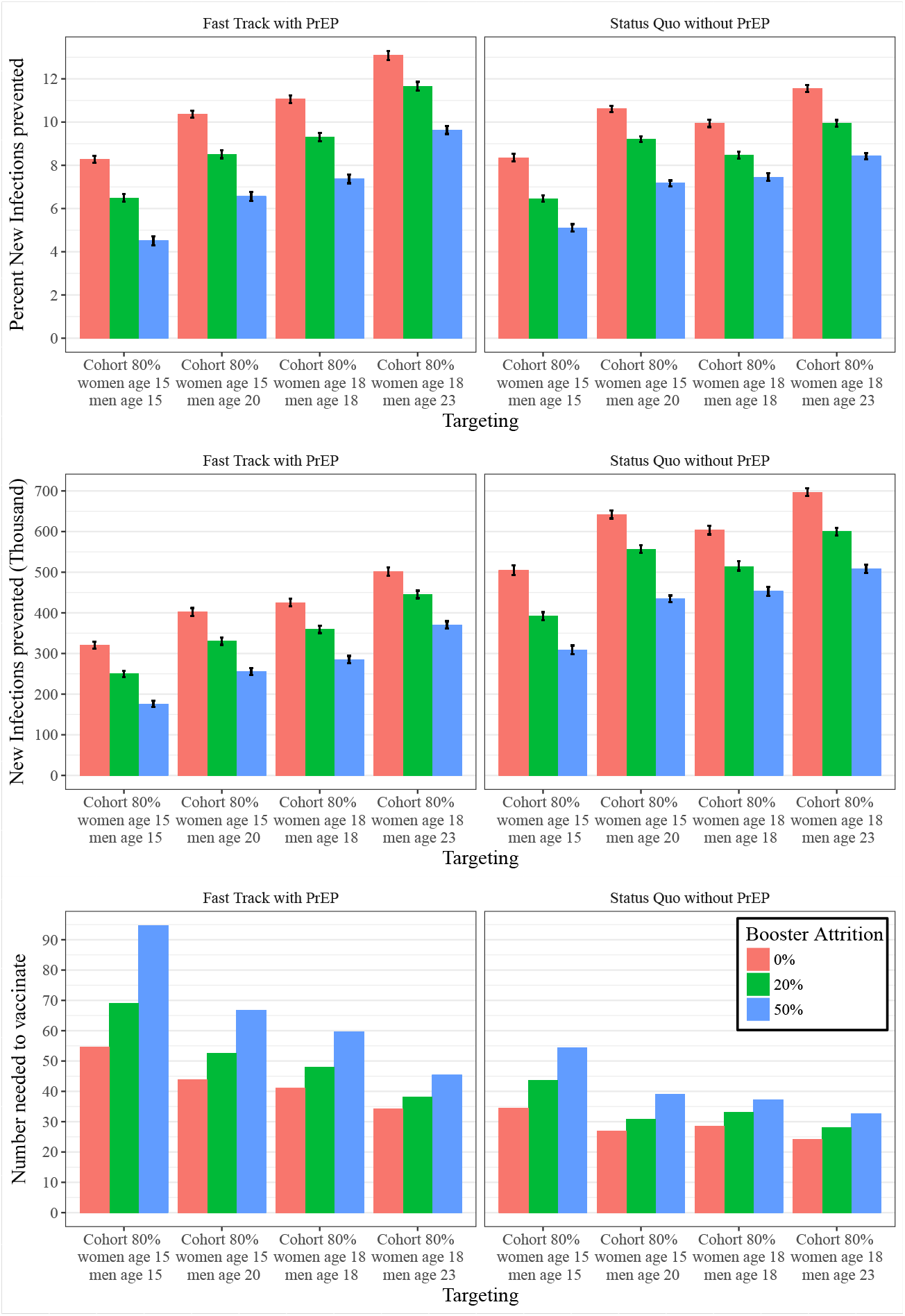
Impact of cohort vaccination at 80% coverage for different treatment scale-up scenarios measured by average number of new infections (first row), percent of new infections prevented (second row), and number needed to vaccinate (third row) between 2027 and 2047 at 50% vaccine efficacy and varying levels of booster attrition (0%, 20% and 50%). Average and 95% confidence intervals are relative to summary statistics across stochastic replicates.

A difference in vaccination age between men and women of 5 years provides additional benefit with up to 402, 000 (95% CI: 392, 000 – 412, 000) and 642, 000 (95% CI: 632, 000 – 652, 000) of new infections prevented for the two scale-up scenarios respectively. Missing 50% of booster would decrease the impact to 256, 000 (95% CI: 247, 000 – 265, 000) and 434, 000 (95% CI: 426, 000 – 443, 000) for vaccination assuming ‘Fast Track with PrEP’ and ‘Status Quo without PrEP’ respectively.

Further impact can be achieved by targeting 18-year-olds resulting in 426, 000 (95% CI: 416,000–435,000) and 604, 000 (95% CI: 593,000–615,000)infections prevented respectively at 0% booster attrition. Echoing the age difference in the peak of HIV incidence, vaccinating 18-year-old women and 23-year-old men would increase the impact to up to 501, 000 (95% CI: 491, 000 – 512, 000) and 697, 000 (95% CI: 687, 000 – 707, 000) infections prevented respectively, given full booster retention.

The impact of age off-setting and vaccination age is also highlighted by the number needed to vaccinate (NNV) to prevent a new infection. When vaccinating at age 15 without versus with age off-set the NNV decreases from 55.7 to 43.9 and from 34.6 to 27.0 respectively. Likewise, vaccinating 18-year-olds requires 41.0 and 28.5 NNV respectively, a five-year age off-setting would decrease these numbers to 34.2 and 24.3 NNV. To illustrate the importance of continued boosting for long-term impact, simulating the latter scenario under 50% (instead of 0%) attrition after each booster dose results in a rise to 45.4 and 32.7 NNV.

Catch-up vaccination appears to be particularly impactful, i.e. by targeting age ranges between 15 to 32 at the beginning of the roll-out at 60% coverage and maintaining cohort coverage of 80% thereafter, the number of new infections prevented increases from 321, 000 to 689, 000 (95%CI: 679, 000–699, 000) (or 18.1% (95%CI:17.9–18.3%)) and from 504, 000 to 941, 000 (95%CI: 932, 000-951, 000) (or 15.6% (95%CI: 15.5 – 15.7%)) for ‘Fast Track with PrEP’ and ‘Status Quo without PrEP’ scenarios respectively (Figure S4). Shifting the age range for catch-up vaccination to 18 to 35 years would have a similar impact of 695, 000 (95%CI: 686, 000 – 704, 000) (or 18.3% (95%CI: 18.1 – 18.5%)) and 935, 000 (95%CI: 924, 000 – 945, 000) (or 15.5% (95%CI: 15.3 – 15.6%)) infections prevented respectively.

Cohort vaccination at 80% or 50% coverage would require approximately 8.4 million or 5.5 million vaccine regimens over 20 years (Table 2). Catchup vaccination would require substantially more vaccine regimens in the early years of the vaccination program, with reduced demand in later years (Figure S4), totaling 15.2 million regimens over 20 years (Table 2). Because of limited durability of vaccine efficacy, the fraction of the sexually active population with partial protection remains small, in spite of high coverage for target populations (Figure S6).

**Table 2:** Economic evaluation

| | Targeting<br>(coverage, gender, age) | Vaccine regimens<sup>a,d</sup><br>(in million) | DALYs averted<sup>a,b</sup><br>(in million) | Vaccine cost<sup>c</sup><br>(in US\$) | ART cost saving<sup>a,b</sup><br>(in million US\$) |
| --- | --- | --- | --- | --- | --- |
| Fast Track with PrEP | 50% men & women age 15 | 5.61 | 0.07 (0-0.16) | 105 (0-236) | 486.33 |
|  | 50% men age 20 & women age 15 | 5.57 | 0.18 (0.1-0.26) | 272 (166-388) | 1158.21 |
|  | 80% men & women age 15 | 8.69 | 0.19 (0.11-0.26) | 176 (103-248) | 1181.60 |
|  | 80% men age 20 & women age 15 | 8.64 | 0.19 (0.11-0.28) | 186 (106-266) | 1255.39 |
|  | 50% men & women age 18 | 5.55 | 0.21 (0.11-0.29) | 307 (187-437) | 1286.80 |
|  | 50% men age 23 & women age 18 | 5.48 | 0.26 (0.16-0.35) | 383 (240-527) | 1592.42 |
|  | 80% men & women age 18 | 8.60 | 0.29 (0.19-0.39) | 277 (193-370) | 1810.46 |
|  | 80% men age 23 & women age 18 | 8.50 | 0.37 (0.27-0.45) | 354 (261-444) | 2286.89 |
|  | 60% men & women age 15-32 | 15.52 | 0.74 (0.65-0.82) | 361 (320-401) | 4543.12 |
|  | 60% men & women age 18-35 | 15.14 | 0.77 (0.69-0.85) | 385 (349-426) | 4705.41 |
| Status Quo without PrEP | 50% men & women age 15 | 5.53 | 0.37 (0.24-0.49) | 533 (352-724) | 2176.25 |
|  | 50% men age 20 & women age 15 | 5.50 | 0.54 (0.41-0.67) | 786 (613-968) | 3192.02 |
|  | 80% men & women age 15 | 8.57 | 0.53 (0.4-0.68) | 500 (371-617) | 3177.77 |
|  | 80% men age 20 & women age 15 | 8.52 | 0.7 (0.56-0.83) | 659 (541-782) | 4158.92 |
|  | 50% men & women age 18 | 5.45 | 0.41 (0.27-0.54) | 604 (394-807) | 2448.40 |
|  | 50% men age 23 & women age 18 | 5.36 | 0.51 (0.36-0.66) | 763 (556-989) | 3022.80 |
|  | 80% men & women age 18 | 8.45 | 0.62 (0.5-0.74) | 589 (481-689) | 3700.01 |
|  | 80% men age 23 & women age 18 | 8.32 | 0.81 (0.68-0.95) | 787 (657-915) | 4833.93 |
|  | 60% men & women age 15-32 | 15.15 | 1.45 (1.32-1.59) | 718 (652-795) | 8634.57 |
|  | 60% men & women age 18-35 | 14.69 | 1.47 (1.33-1.6) | 749 (679-810) | 8665.38 |

All numbers are averaged over 50 simulations, with full booster retention. 95% confidence intervals, if provided, are in parentheses.
<sup>a</sup> Cumulative sum 2027-2047
<sup>b</sup> 5% annual discount starting in 2018
<sup>c</sup> Maximum vaccine cost at a cost-effectiveness threshold of 1x GDP. This takes ART cost saving into account, it is not an additional benefit.
<sup>d</sup> Number of primary series of five vaccinations administered

### Vaccination Cost Thresholds

For economic evaluation of vaccine cost per regimen (including delivery and implementation costs), we calculated the maximum cost at which vaccination would remain cost-effective, referred to as maximum vaccine cost in what follows. Cost-effectiveness thresholds are relative to the 2015 gross domestic product (GDP) of South Africa or $5, 691 USD per capita [24]. Since future HIV incidence and booster attrition rates remain highly speculative, cost and DALYs estimates are averaged across simulations with 0% booster attrition, such that the resulting vaccine cost is an upper bound in terms of durability. For cohort vaccination at 80% coverage with a cost-effectiveness threshold of 1x per-capita GDP per DALY averted, the average maximum vaccine cost (for both vaccine product and implementation of a 10-year vaccine series) ranges from $176 (95% CI: $103 – $248) to $354 (95% CI: $261-$444) (Table 2) in ‘Fast Track with PrEP’ scenarios. Under the same scale-up assumption, at 60% catch-up vaccination, maximum cost-effective vaccine cost ranges from $361 (95% CI: $320-$401) to $385 (95% CI: $349-$426). Although the catch-up strategy requires substantially more vaccine regimens, it also offers higher ART cost savings (Table 2). For the ‘Status Quo without PrEP’ scenario, maximum vaccine cost would more than double, ranging between $500 (95% CI: $371-$617) and $749 (95% CI: $679 – $810) for cohort and catch-up vaccination respectively at 80% coverage. Lowering the annual discount rate to emphasize present value to society [44] would increase estimated maximum vaccine costs (Figure S7) from 176 US$ and 385 US$ to up to $480 and $548 for cohort at 80% coverage and catch-up vaccination at 60% coverage respectively under ‘Fast Track with PrEP’ assumptions.

Among the simulated vaccination strategies, catch-up vaccination age 18-35 is the most efficient when compared to cohort vaccination, since ART cost savings as well as averted DALYs are high, despite very similar maximum vaccine cost. Sensitivity analyses shows that maximum vaccine cost is proportional to the CET, e.g. at a threshold of approximately 750$ per DALY averted (see Discussion for details), maximum vaccine cost would range between $22 and $60 under ‘Fast Track with PrEP’ assumptions and $74 and $115 in the ‘Status Quo without PrEP’ scenario (Figure S8).

## 4. Discussion

The results of this study highlight the need to exploit different vaccination targeting and roll-out scenarios to maximize the population-level impact of a partially effective HIV vaccine in South Africa. Even under high attrition rates, providing additional booster doses for up to one decade after the primary series can increase public health impact, as long as gender-specific age ranges of highest incidence are covered by vaccination. Although adolescent HIV vaccination has the potential of reaching high coverage in South Africa through school-based programs (such as Human Papilloma Virus routine vaccination [45]), our modeling results (see Table 2 and Table S3) do not make a strong case for vaccinating before the age of sexual debut, estimated at a median of 18 years of age [46]. The same conclusion applies to the potential impact on vertical transmission by vaccinating 15-year-old women. Prevention of mother-to-child transmission is efficacious in South Africa (only 1.3 % of live births in 2017 were HIV positive [47]) and only 12% of live births are given at age 19 or younger whereas 70% of births are given at ages 20-34 [48].

The time-dependent course of vaccine efficacy necessitates aligning vaccination such that the times of highest efficacy are within the ages of highest HIV incidence in 2027 – projected to be 20-25 years of age for women, and five to ten years older for men (Figure S9). Specifically, our results indicate that, of the gender/age combinations we compared, the greatest public health impact of the pox-protein HIV vaccine would be achieved by vaccinating 18-year-old women and 23-year-old men.

Long-term economic evaluation of vaccination suffers, in part, from the uncertainty of whether HIV treatment and prevention scale-up could be achieved between now and 2027. We derived estimates for the maximum vaccine cost (10-year regimen, including product and delivery costs) to remain cost-effective averaging for two different treatment and prevention scale-up projections. Our analysis suggested an estimated maximum cost of $105 – $787 at which vaccination with a full 10-year regimen would be cost-effective based on a 1xGDP per capita threshold [49]. Measuring opportunity cost from a government perspective and deriving cost-effectiveness thresholds for health care interventions for particular countries is an area of active research and debate [50]. Instead of using the WHO cost-effectiveness thresholds, the South African HIV/TB investment case study group developed its own CE threshold in determining the cost-effectiveness of new HIV interventions in South Africa. Using an iterative optimization approach for all existing treatment and prevention options, the group concluded that a cost-effectiveness threshold of approximately $750 per life-year saved would be an appropriate upper bound for the incremental cost to South Africa’s HIV program compared to the baseline scenario (maintained coverage of all interventions at 2014 levels). This reflects the fact that HIV treatment and prevention uptake is already saturated in South Africa, and suggests that only a limited number of interventions should be scaled up to avoid detrimental population health outcomes resulting from budget funds being displaced from other interventions.

There are several major limitations to this analysis. The time-dependent vaccine efficacy curves are based on limited data from RV144 and goals for the pox-protein vaccine regimen, and we did not consider any primary series attrition. Additional sources of uncertainty include differences in regimens and study populations between RV144 and the ongoing trial, statistical uncertainty in the RV144 results, and translation of cohort efficacy data to an individual-based model in the absence of HIV exposure information for the cohort under study. We did not model the impact of possible changes in voluntary medical male circumcision scale-up on vaccination, nor did we model the recently added 18-month dose in the amended trial protocol. Since available data on the distribution of earnings, non-healthcare consumption costs, and patient time costs in South Africa were insufficient to be included in this analysis, we adopted a government payer instead of a societal perspective. The economic evaluation of the vaccine does not account for indirect economic implications (e.g. productivity effects). The threshold analysis to determine the maximum cost-effective cost per vaccine regimen included product, delivery and implementation cost, which may vary across the simulated strategies, making cost comparisons difficult. Although the estimated maximum vaccine cost accounts for possible resource displacement (e.g. ART cost saving), further additional investment might be necessary to maintain treatment and prevention programs with concurrent vaccine roll-out, such that vaccine cost is likely underestimated. The lack of bounds on the vaccine price and insufficient data on other cost-drivers such as staff training, information campaigns and monitoring for such a complex regimen made it challenging to carry out a well-informed cost breakdown analysis. Furthermore, different target populations may be more or less challenging to reach. This is difficult to know a priori and was not included in the analysis, and therefore remains an area important to explore in close collaboration with implementers of HIV prevention programs. We did not model risk compensation [15, 51] i.e. increases in risky behavior by vaccine recipients, bearing in mind that a partially effective vaccine might necessitate additional counseling to prevent the false impression of full protection. Vaccine-induced seropositivity [52] is likely to add substantial additional cost [53] due to the need to use nucleic acid based HIV testing to distinguish HIV infections from the presence of vaccine-induced HIV antibodies. Finally, we likely underestimate the long-term benefits of vaccination. DALY calculations were truncated at the vaccination endpoint in 2047, i.e. vaccination could add at most 20 additional years of life from the start of vaccination to the endpoint.

## 5. Conclusion

Taken together, our model suggests that, averaged across different treatment scale-up settings, vaccinating population cohorts aligned with the ages of highest HIV incidence (i.e. 18-year-old women and 23-year-old men) including continued boosting for up to 10 years could avert up to 13.1% and 11.5% of HIV infections in the coming decades (for ‘Fast Track with PrEP’ and ‘Status Quo without PrEP’ respectively). If durability of vaccine efficacy proves to be better than observed in the RV144 trial, the benefit could be appreciably larger. Adult catch-up vaccination and efforts to ensure continued boosting, could further increase the impact of the vaccination to possibly just over 18.3% (or 694, 000) and 15.5% (or 935, 000) of new infections prevented (for ‘Fast Track with PrEP’ and ‘Status Quo without PrEP’ respectively). However, we recognize that identifying optimal vaccine implementation platforms and deployment channels to deliver this complex vaccine regimen at high coverage will be a significant challenge–especially when vaccinating 18 year-old women instead of adolescent girls who may be more easily reached through a school-based program. Therefore, improving deployability of HIV vaccine regimen by optimizing vaccine performance characteristics (e.g. increased vaccine efficacy, longer duration of protection, fewer required doses) should be ultimate goal of future vaccine research. Hopefully, such optimization would be supported by the identification of immune correlates of protection in HVTN 702. Although the maximal impact of 18.3% of new infection prevented is appreciable, the rollout of a partially effective, rapidly waning vaccine alone will not eliminate HIV as a public health priority in South Africa. Therefore, vaccination should be performed in parallel with continued innovation in HIV prevention technologies.

## Acknowledgements

We acknowledge productive discussions with Dan Klein, Dan Bridenbecker, and Mandy Izzo (all Institute for Disease Modeling), Peter Gilbert (Fred Hutchinson Cancer Research Center), Leigh Johnson (University of Cape Town), the Vaccine Research Program at the National Institutes of Allergy and Infectious Disease (US National Institutes of Health), members of the P5 Global Access Committee (comprised of representatives from the Bill & Melinda Gates Foundation, National Institutes of Health (DAIDS/NIAID), Sanofi Pasteur, Glaxo-SmithKline and the South African Medical Research Council) and Shift Health (Toronto, Canada) and the HIV Modelling Consortium which facilitated the Modelling workshop convened by the P5 GAC. This work was supported by Bill and Melinda Gates through the Global Good Fund.

## Author contributions

C.S., A.B., D.T.D., T.B.H., A.N.P. conceived the study and designed the experiments. C.S. performed the modeling experiments. C.S., A.B., D.T.D., B.J.A., P.R., T.B.H., A.N.P., L.-G. B., H.R., G.G. interpreted the data and contributed to writing the manuscript.

## Conflict of interest

The authors declare no competing interests.

Appendix A. Supplementary Material

