## Supplementary Material for "Targeting and Vaccine Durability Are Key for Population-level Impact and Cost-Effectiveness of a Pox-Protein HIV Vaccine Regimen in South Africa"

---

*Keywords:* HIV vaccine, South Africa, agent-based modeling, cost-effectiveness

---

**Model set-up and calibration**. EMOD-HIV is an individual-based model that simulates transmission of HIV using an explicitly defined network of heterosexual relationships that are formed and dissolved according to age- and risk-dependent preference patterns [1]. The synthetic population was initiated in 1960, and population recruitment and mortality was assumed to be proportional following age- and gender-stratified fertility and mortality tables and projections from the 2012 UN World Population Prospects [2]. Since the population size of South Africa exceeds the computational limit of simulated agents, we assumed that one simulated agent corresponds to 300 real-world individuals. The model was calibrated to match retrospective estimates of age- and gender-stratified, national-level prevalence, incidence, and ART coverage from four nationally representative HIV surveys in South Africa [3, 4, 5, 6, 7]. For each simulated vaccine scenario, we used the 50 most likely parameter sets. For the purpose of this modeling study, we emphasized calibration of model parameters concerning risk assortativity during partnership formation, duration and concurrency of partnerships as well as condom usage to fit gender- and age-stratified incidence and prevalence data (Figure S1). The age patterns of sexual mixing were configured to match those observed in the rural, HIV-hyperendemic province of KwaZulu-Natal, South Africa [8, 1]. Recently, a validation study showed that self-reported partner ages in this setting are relatively accurate, with 72% of self-reported estimates falling within two years of the partners actual date of birth [9]. Further, the transmission patterns observed in EMOD [10] are consistent with those revealed in a recent phylogenetic analysis of the age/gender patterns of HIV transmission in this setting [11]. The EMOD model also includes vertical transmission from pregnant mothers to children.

Transmission rates within relationships depend on HIV disease stage, male circumcision, condom usage, co-infections. Viral suppression achieved through antiretroviral therapy [12, 13] is assumed to reduce transmission by 92%-an estimate based on observational data of serodiscordant couples in which outside partnerships could have contributed to HIV acquisition [14].

**HIV Treatment and Prevention**. We configured the EMOD health care system module to follow trends in antiretroviral therapy (ART) expansion in South Africa. Treatment begins with voluntary counseling and

*Corresponding author

testing (VCT), antenatal and infant testing, symptom-driven testing, and low level of couples testing. The model includes loss to follow-up between diagnosis and staging, between staging and linkage to ART or pre-ART care, and during ART or pre-ART care [15]. Projections of South Africa treatment expansion in the no vaccine reference group are calibrated to reflect a gradual decline of HIV incidence without elimination, so that HIV remains endemic through 2050 [16]. All scenarios included medical male circumcision [17] at 22% coverage and conferring 60% reduction in acquisition risk with lifetime durability. Condom usage was dependent on four relationship types (transitory, informal, marital, commercial), with per act usage probability ramping up to median values of 62%, 39%, 26%, and 85% by 2027 across parameter draws. The commercial relationship type was implemented to model partnership dynamics between sex workers and patrons with high concurrency and short relationship duration. Based on adherence patterns and efficacy results from the Partners PrEP trial [18], we assumed that oral pre-exposure prophylaxis (PrEP) had 74% efficacy in terms of risk reduction for young adults.

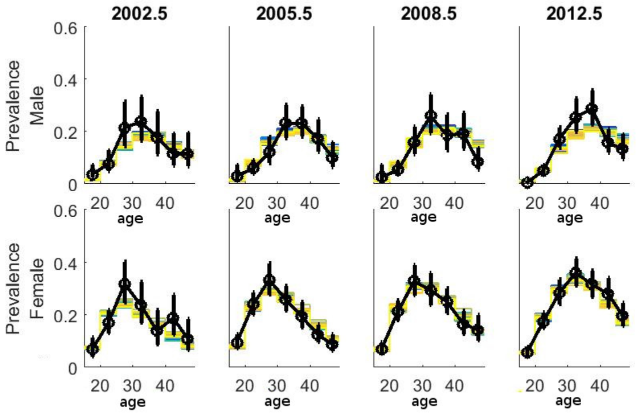


(a) Prevalence stratified by age (15-24, 25-29, 30-34, 35-39, 40-44, 45-49) and gender compared to HSRC survey data from 2002, 2005, 2008 and 2012.

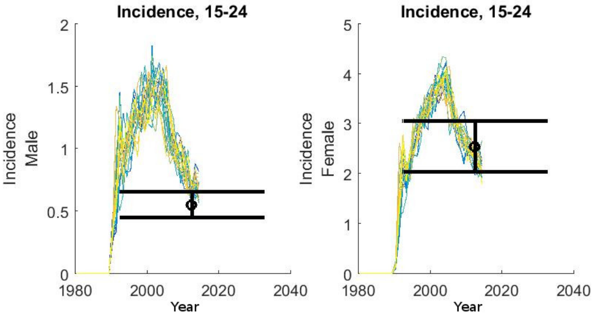


(b) Incidence age 15-24, stratified by gender

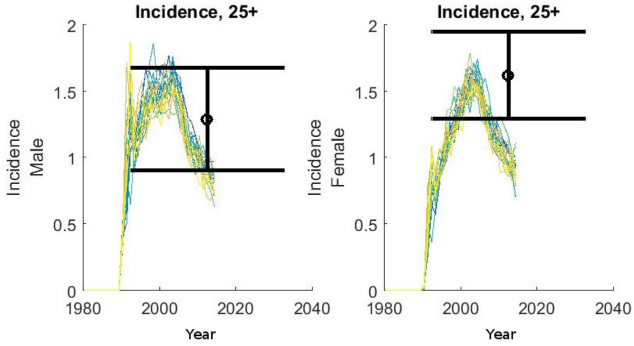


(c) Incidence age 25 and older, stratified by gender

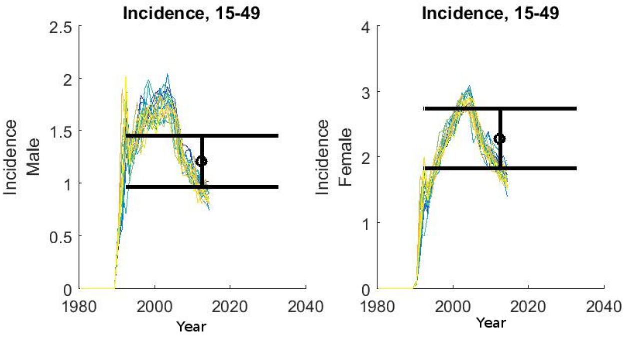


(d) Incidence age 15-49, stratified by gender

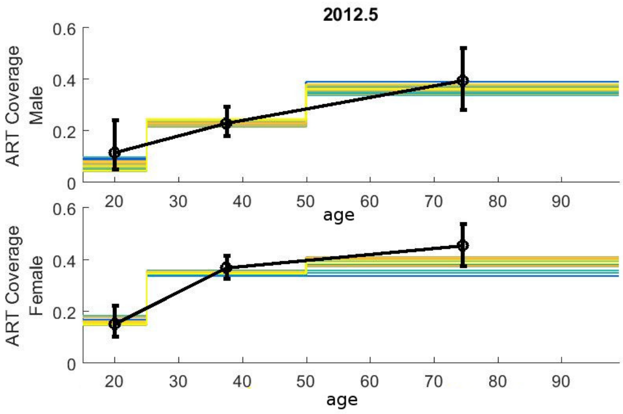


(e) ART coverage, stratified by age (15-24, 25-29, 30-34, 35-39, 40-44, 45-49) and gender

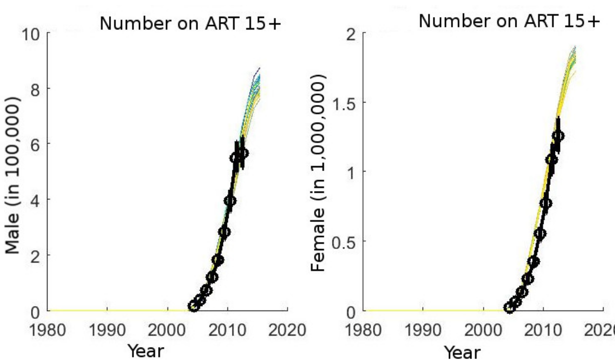


(f) Number on antiretroviral therapy (ART) stratified by gender

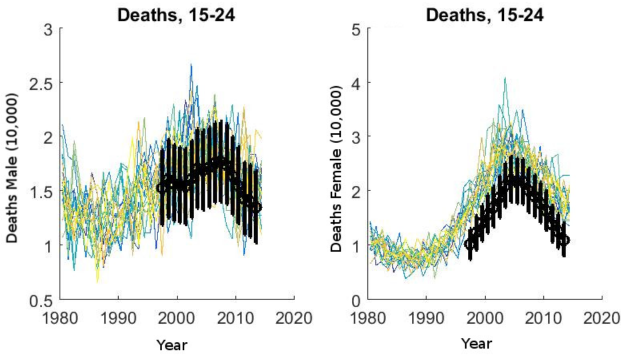


(g) Deaths (all cause mortality) by gender for age 15-24.

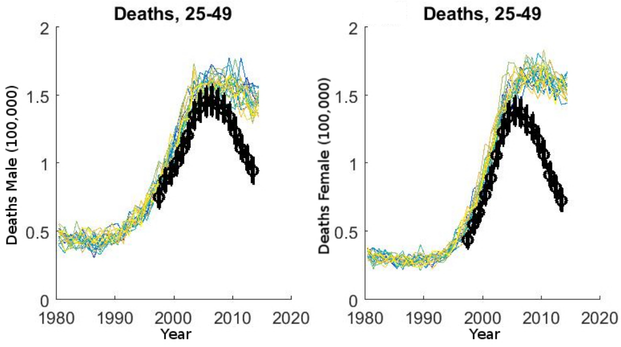


(h) Deaths (all cause mortality) by gender for age 25-49.

Figure S1: Results of model calibration (trajectories in colors) to incidence, prevalence, ART coverage and all-cause mortality estimates [3, 4, 5, 6, 7, 19] for South Africa (black).

Table S1: Cohort vaccination scenarios, targeting individuals of specific ages every year. School-age is defined by receiving the primary vaccination series before the age of 18, age off-set refers to a difference in target age of five years between men and women.

| Cohort scenario | male | | female | |
|---|---|---|---|---|
| | age | coverage | age | coverage |
| 1 | 10 | 30% | 10 | 30% |
| 2 | 10 | 50% | 10 | 50% |
| 3 | 10 | 80% | 10 | 80% |
| 4 | 10 | 0% | 10 | 80% |
| 5 | 10 | 0% | 10 | 80% |
| 6 | 10 | 0% | 10 | 80% |
| 7 | 15 | 30% | 15 | 30% |
| 8 | 15 | 50% | 15 | 50% |
| 9 | 15 | 80% | 15 | 80% |
| 10 | 20 | 30% | 15 | 30% |
| 11 | 20 | 50% | 15 | 50% |
| 12 | 20 | 80% | 15 | 80% |
| 13 | 0 | 30% | 15 | 30% |
| 14 | 0 | 50% | 15 | 50% |
| 15 | 0 | 80% | 15 | 80% |
| 16 | 18 | 30% | 18 | 30% |
| 17 | 18 | 50% | 18 | 50% |
| 18 | 18 | 80% | 18 | 80% |
| 19 | 23 | 30% | 18 | 30% |
| 20 | 23 | 50% | 18 | 50% |
| 21 | 23 | 80% | 18 | 80% |
| 22 | 18 | 0% | 18 | 80% |
| 23 | 18 | 0% | 18 | 80% |
| 24 | 18 | 0% | 18 | 80% |

Table S2: Catch-up vaccination, targeting 2 different age ranges for 5 years, followed by cohort vaccination.

| Catch-up scenario | age range (male & female) | ramp-up coverage | mainte-nance coverage |
|---|---|---|---|
| 1 | 15-32 | 10% | 30% |
| 2 | 15-32 | 30% | 50% |
| 3 | 15-32 | 60% | 80% |
| 4 | 18-35 | 10% | 30% |
| 5 | 18-35 | 30% | 50% |
| 6 | 18-35 | 60% | 80% |

Table S3: Economic evaluation

| | Targeting (coverage, gender, age) | Vaccine regimens[a,d] (in million) | DALYs averted[a,b] (in million) | Vaccine cost[c] (in US$) | ART cost saving[a,b] (in million US$) |
|---|---|---|---|---|---|
| Fast Track with PrEP | 30% men & women age 10 | 3.56 | 0 (0-0.07) | 0 (0-153) | $< 0$ |
| | 50% men & women age 10 | 5.81 | 0 (0-0.04) | 0 (0-54) | $< 0$ |
| | 50% women age 10 | 2.92 | 0 (0-0.07) | 0 (0-236) | $< 0$ |
| | 80% men & women age 10 | 8.89 | 0 (0-0.05) | 0 (0-40) | $< 0$ |
| | 80% women age 10 | 4.47 | 0.01 (0-0.11) | 26 (0-213) | 96.84 |
| | 30% men & women age 15 | 3.42 | 0.07 (0-0.16) | 169 (0-365) | 460.55 |
| | 30% men age 15 & women age 20 | 3.40 | 0.09 (0.01-0.18) | 216 (21-413) | 582.52 |
| | 50% women age 15 | 2.82 | 0.09 (0.01-0.18) | 277 (30-502) | 605.99 |
| | 80% women age 15 | 4.37 | 0.16 (0.09-0.23) | 308 (184-433) | 1046.11 |
| | 10% men & women age 15-32 | 3.66 | 0.03 (0-0.11) | 69 (0-238) | 213.89 |
| | 30% men & women age 15-32 | 7.96 | 0.36 (0.28-0.44) | 347 (273-416) | 2191.12 |
| | 30% men & women age 18 | 3.39 | 0.07 (0-0.19) | 164 (0-441) | 437.26 |
| | 30% men age 18 & women age 23 | 3.34 | 0.17 (0.09-0.24) | 409 (229-596) | 1039.51 |
| | 50% women age 18 | 2.76 | 0.13 (0.04-0.22) | 391 (128-665) | 828.07 |
| | 80% women age 18 | 4.29 | 0.24 (0.15-0.33) | 460 (302-624) | 1506.84 |
| | 10% men & women age 18-35 | 3.59 | 0.11 (0.03-0.18) | 242 (52-420) | 668.37 |
| | 30% men & women age 18-35 | 7.79 | 0.38 (0.29-0.46) | 373 (286-461) | 2297.36 |
| Status Quo without PrEP | 30% men & women age 10 | 3.48 | 0.07 (0-0.22) | 172 (0-498) | 431.40 |
| | 50% men & women age 10 | 5.69 | 0.04 (0-0.17) | 55 (0-241) | 230.80 |
| | 50% women age 10 | 2.86 | 0.07 (0-0.21) | 207 (0-562) | 423.23 |
| | 80% men & women age 10 | 8.71 | 0.02 (0-0.14) | 18 (0-121) | 112.72 |
| | 80% women age 10 | 4.38 | 0.04 (0-0.16) | 72 (0-272) | 224.32 |
| | 30% men & women age 15 | 3.37 | 0.21 (0.1-0.32) | 501 (246-731) | 1264.78 |
| | 30% men age 15 & women age 20 | 3.35 | 0.29 (0.17-0.41) | 693 (404-943) | 1723.22 |
| | 50% women age 15 | 2.78 | 0.26 (0.14-0.37) | 758 (420-1089) | 1568.14 |
| | 80% women age 15 | 4.31 | 0.44 (0.31-0.56) | 810 (589-1051) | 2579.00 |
| | 10% men & women age 15-32 | 3.58 | 0.31 (0.18-0.45) | 684 (388-994) | 1808.20 |
| | 30% men & women age 15-32 | 7.78 | 0.76 (0.61-0.91) | 746 (607-882) | 4495.13 |
| | 30% men & women age 18 | 3.32 | 0.25 (0.11-0.39) | 599 (273-929) | 1480.66 |
| | 30% men age 18 & women age 23 | 3.27 | 0.4 (0.24-0.55) | 973 (589-1330) | 2331.69 |
| | 50% women age 18 | 2.70 | 0.3 (0.15-0.46) | 903 (452-1316) | 1815.89 |
| | 80% women age 18 | 4.18 | 0.43 (0.3-0.57) | 834 (580-1090) | 2600.30 |
| | 10% men & women age 18-35 | 3.49 | 0.32 (0.17-0.47) | 717 (379-1048) | 1837.07 |
| | 30% men & women age 18-35 | 7.57 | 0.71 (0.58-0.86) | 723 (589-859) | 4223.91 |

All numbers are averaged over 50 simulations, with full booster retention. 95% confidence intervals, if provided, are in parentheses.

[a] Cumulative sum 2027-2047

[b] 5% annual discount starting in 2018

[c] Maximum vaccine cost at a cost-effectiveness threshold of 1x GDP. This takes ART cost saving into account, it is not an additional benefit.

[d] Number of primary series of five vaccinations administered

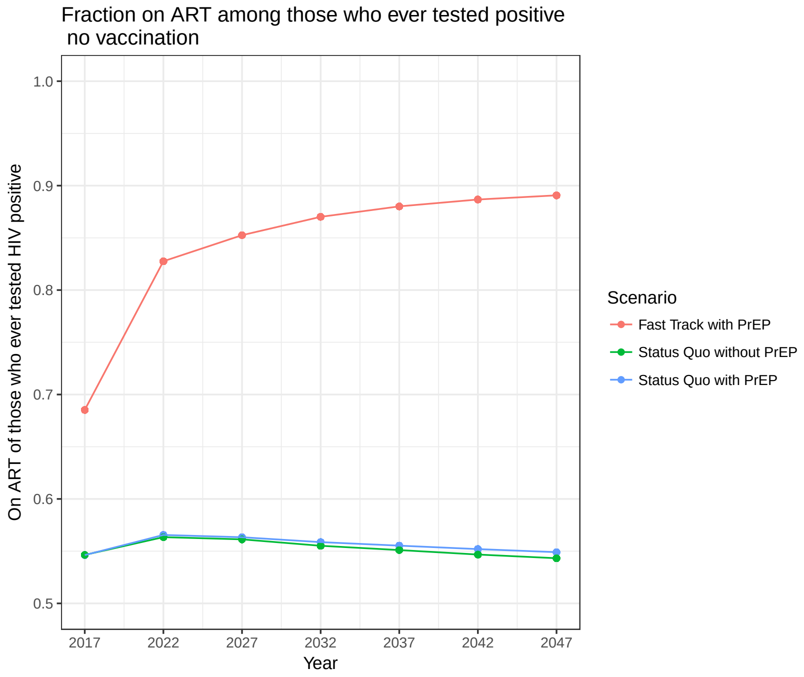


Figure S2: ART coverage for individuals who tested HIV positive for different scale-up scenarios.

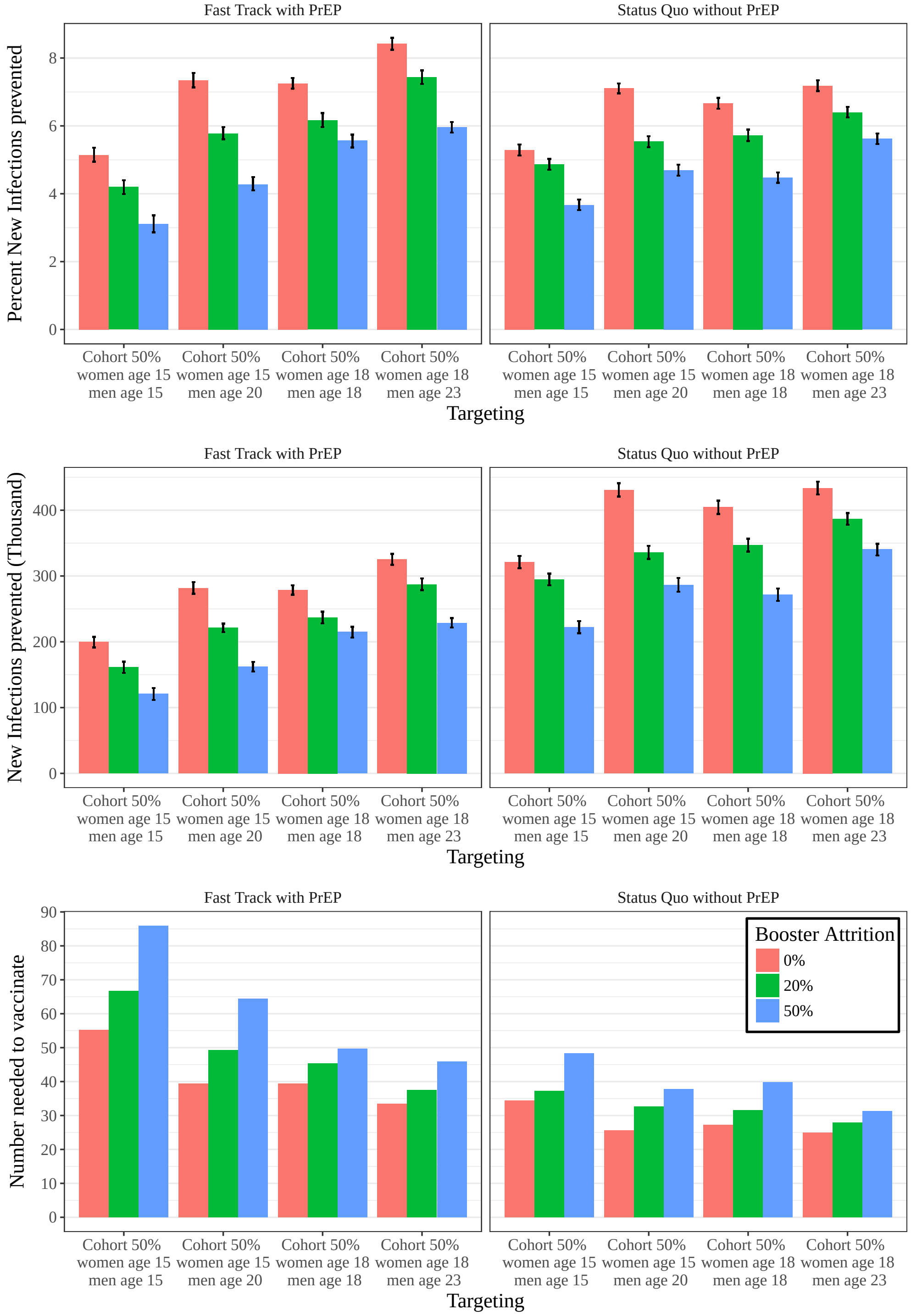


Figure S3: Impact of cohort vaccination at 50% coverage for different treatment scale-up scenarios measured by (A) average number of new infections, (B) percent of new infections prevented, and (C) number needed to vaccinate (NNV) between 2027 and 2047 at 50% vaccine efficacy and varying levels of booster attrition (0%, 20% and 50%). Average and 95% confidence intervals are relative to summary statistics across stochastic replicates.

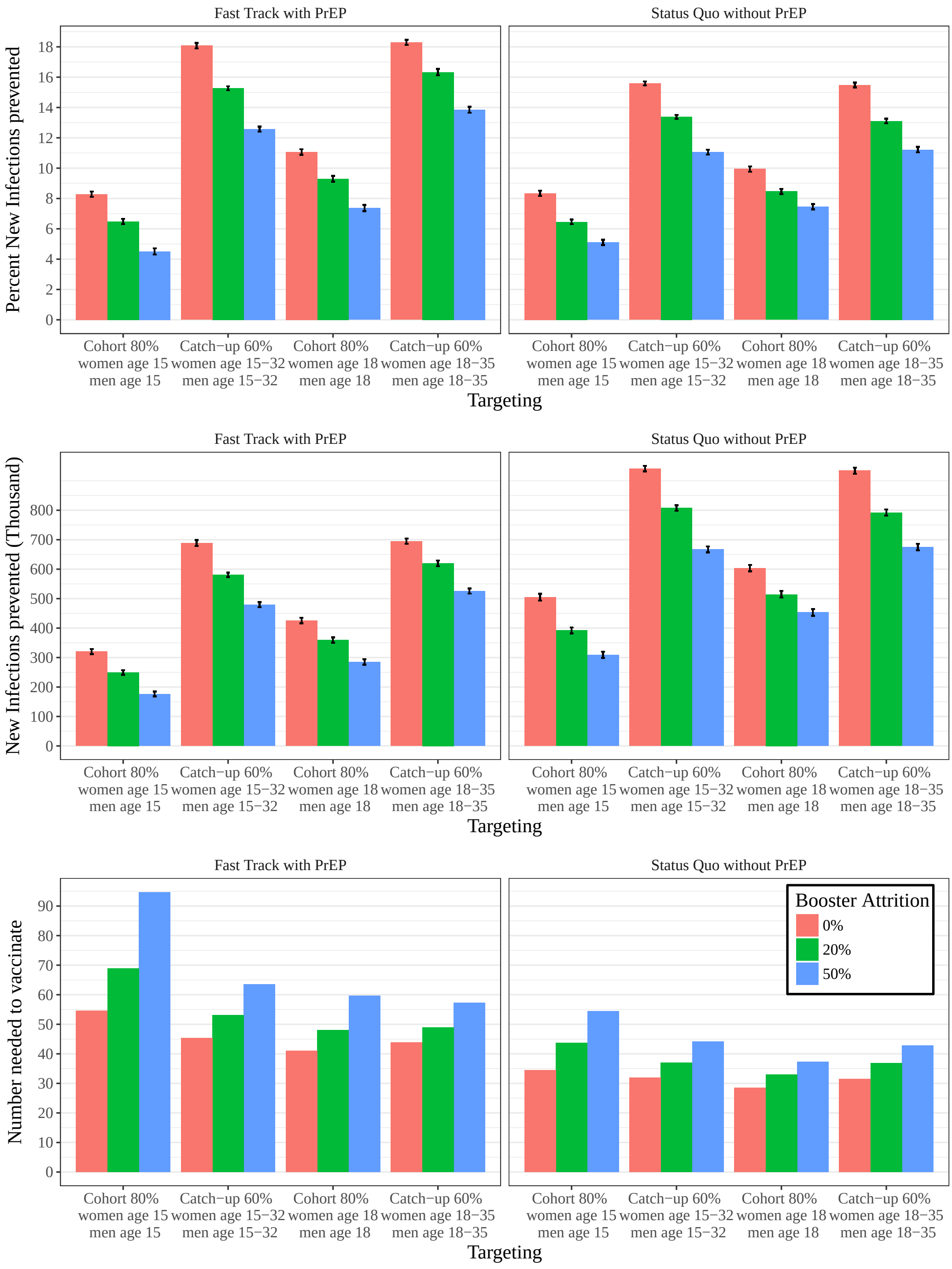


Figure S4: Impact of cohort vs catch-up vaccination at 80% coverage for different treatment scale-up scenarios measured by (A) average number of new infections, (B) percent of new infections prevented, and (C) number needed to vaccinate (NNV) between 2027 and 2047 at 50% vaccine efficacy and varying levels of booster attrition (0%, 20% and 50%). Average and 95% confidence intervals are relative to summary statistics across stochastic replicates.

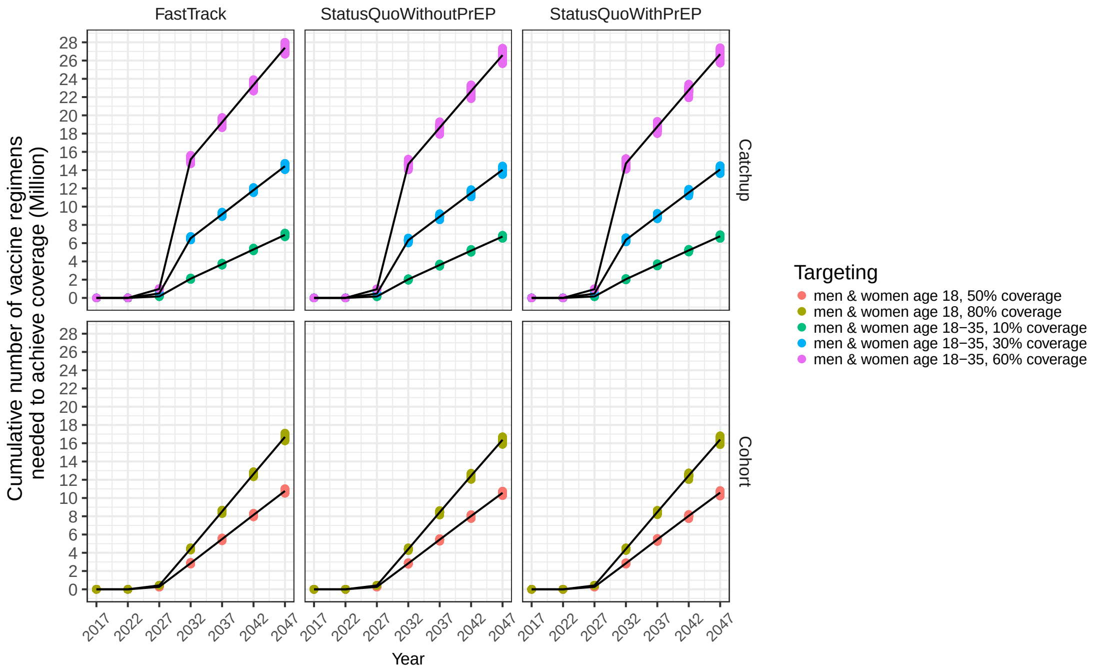


Figure S5: Number of vaccine regimens needed to achieve coverage.

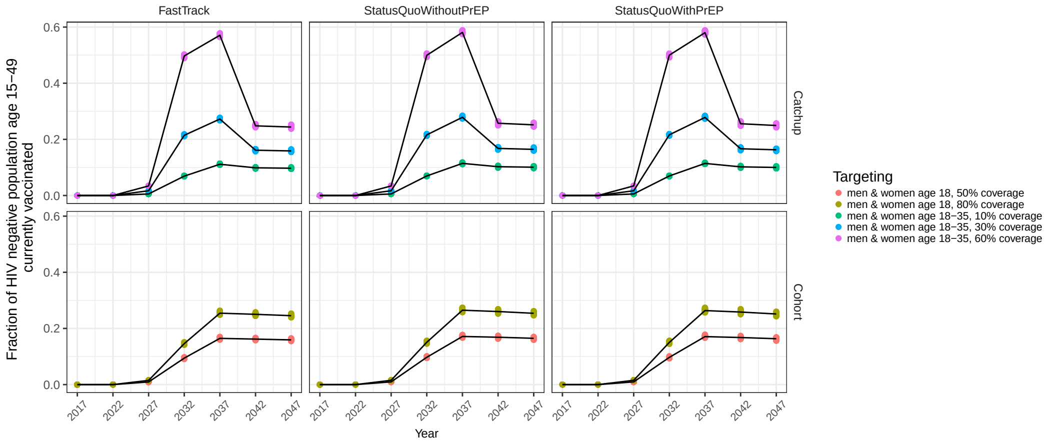


Figure S6: Fraction of HIV negative population age 15-49 currently vaccinated.

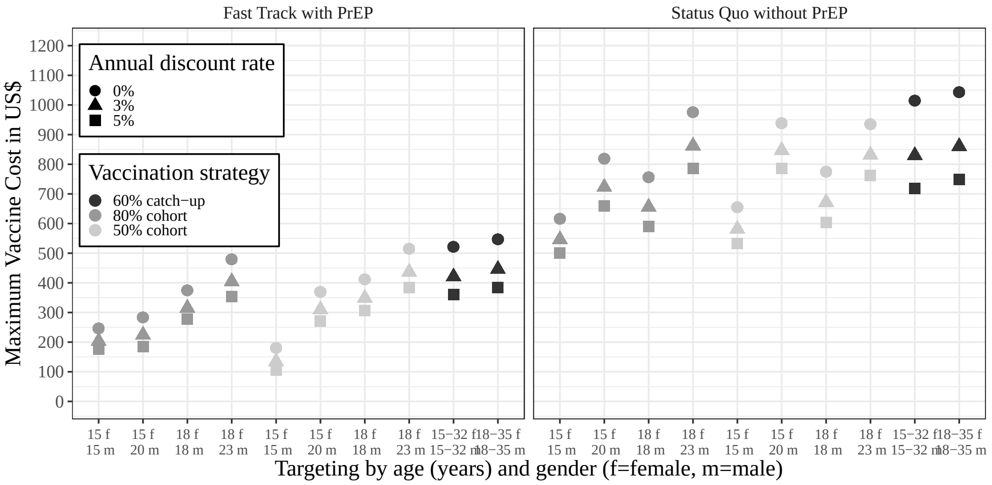


Figure S7: Maximum vaccine cost by targeting strategies and treatment scale-up assumption, at full booster retention and annual discount rates of 0, 3 and 5%.

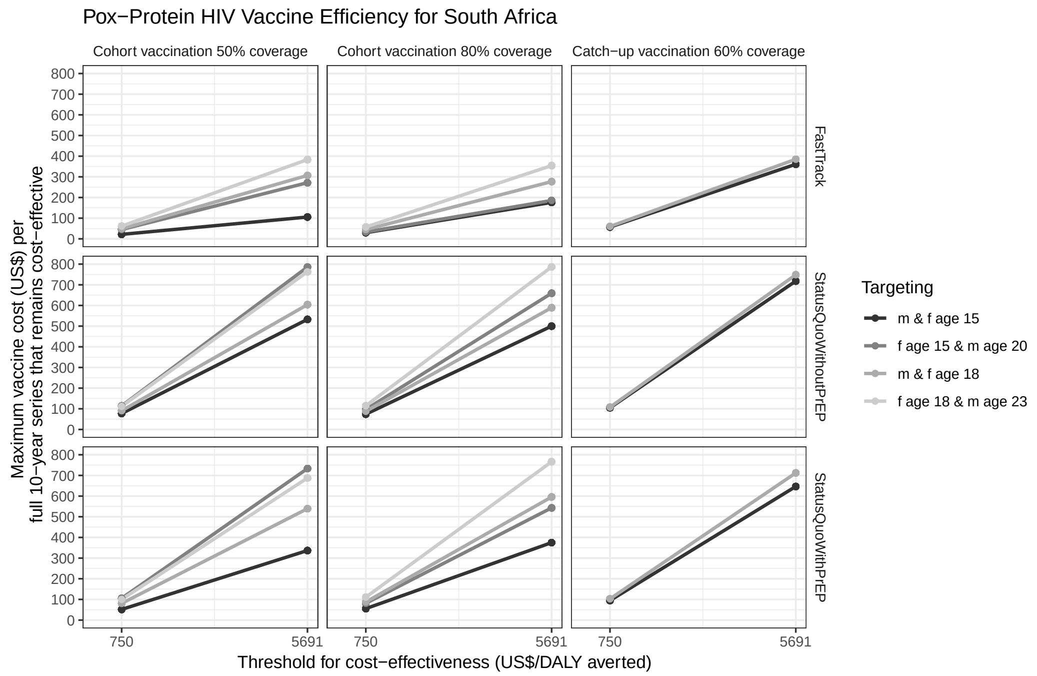


Figure S8: Vaccine efficiency evaluated in terms of maximum vaccine cost that remains cost-effective. Cost-effectiveness thresholds were defined as 750 and 5691 US$ per capita per DALY averted. We consider vaccine cost for a full 10-year series of vaccination including delivery. For cost evaluation, DALYs and all cost were discounted at 5% annual rate. All reported values were averaged across stochastic replicates, at a 0% booster attrition level.

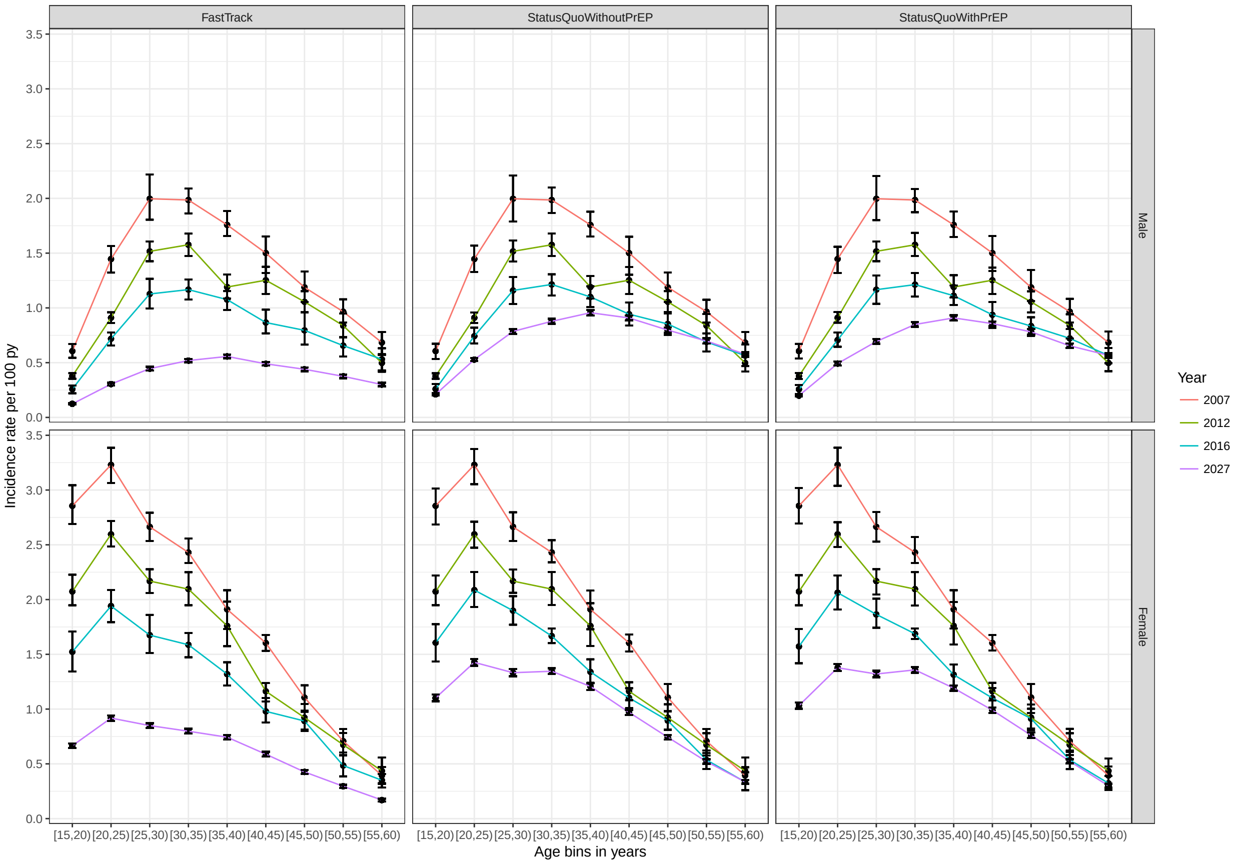


Figure S9: Age-stratified incidence in 2007, 2012, 2016 and 2027 averaged for three different scale-up scenarios.
